# Ecological and Evolutionary Oscillations in Host-Parasite Population Dynamics, and The Red Queen

**DOI:** 10.1101/001735

**Authors:** Jomar F. Rabajante

**Affiliations:** Institute of Mathematical Sciences and Physics, University of the Philippines Los Baños, Laguna, Philippines; and Graduate School of Science and Technology, Shizuoka University, Japan

**Keywords:** host-parasite, prey-predator, coevolution, oscillation, Red Queen, selection gradient

## Abstract

In a host-parasite system, the constitutive interaction among the species, regulated by the growth rates and functional response, may induce populations to approach equilibrium or sometimes to exhibit simple cycles or peculiar oscillations, such as chaos. A large carrying capacity coupled with appropriate parasitism effectiveness frequently drives long-term apparent oscillatory dynamics in population size. We name these oscillations due to the structure of the constitutive interaction among species as *ecological*.

On the other hand, there are also exceptional cases when the evolving quantitative traits of the hosts and parasites induce oscillating population size, which we call as *evolutionary*. This oscillatory behavior is dependent on the speed of evolutionary adaptation and degree of evolutionary trade-off. A moderate level of negative trade-off is essential for the existence of oscillations. Evolutionary oscillations due to the host-parasite coevolution (known as the Red Queen) can be observed beyond the ecological oscillations, especially when there are more than two competing species involved.

**One Sentence Summary:** We investigate several cases yielding to oscillating host-parasite populations, and we found that the Red Queen hypothesis can explain some of the exceptional cases.

Graphical Abstract:

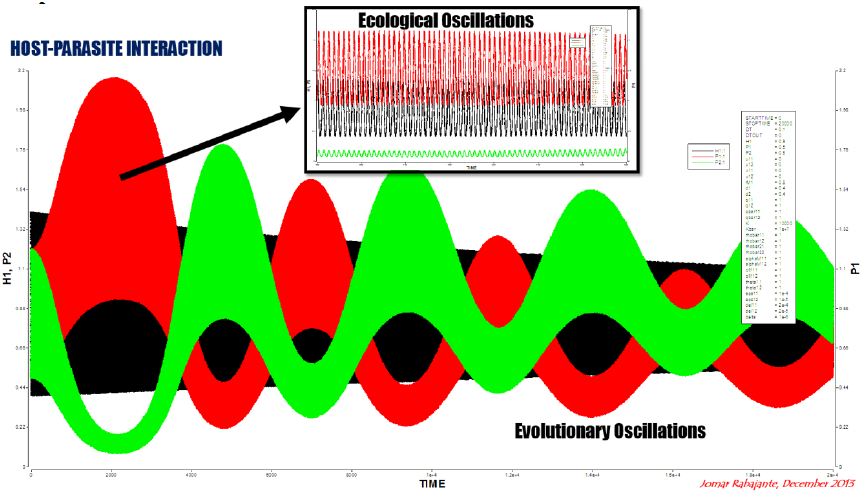

## I. Introduction

Interacting species can coexist but it is possible that some species approach the edge of extinction. In most cases, the species with the greatest fitness are always dominant, but there are also cases when populations exhibit fluctuating population size. In this paper, we investigate host-parasite population dynamics using an ordinary differential equation (ODE) model to particularly determine several cases yielding to long-term apparent oscillatory behavior. Exploring the realm of fluctuating, and possibly non-equilibrium, dynamics can help explain complex antagonistic systems. The abstract model of host-parasite interaction can also describe competition such as predator-prey and exploiter-victim.

Various researches have been done analyzing and employing multispecies host-parasite models to represent theoretical, experimental and natural situations [Jost, 1998; Briggs and Hoopes, 2004; Piana et al., 2006; Beninca et al., 2009; Mougi and Kondoh, 2012]. Solution to a model may converge to an equilibrium point, which is either a stable node or stable focus. In some cases, solution may approach a limit cycle, which is possibly synchronous or asynchronous. There are also situations where fluctuating behavior leads to chaos. Oscillating population size (e.g., solution converging to a stable focus, limit cycle or strange attractor) is not unusual in nature, and in fact, is very important in maintaining diversity. This may be a result of the average antagonistic interaction among species (deterministic) or a result of randomness (stochastic). Time delays and environmental factors can also play significant roles in producing oscillations.

An interaction model can involve single host and single parasite; many hosts but single parasite; single host but many parasites; or many hosts and many parasites. ODE models can differ on the growth terms (e.g., exponential growth or logistic growth of host species), and functional response (e.g., Holling type I, II or III, Beddington-DeAngelis and Monod-Haldane). Analysis of classical and low-dimensional host-parasite interaction models, such the Lotka-Volterra (LV) and Rosenzweig-MacArthur (RM), can be found in various literatures [Korobeinikov and Wake, 1999; Murray, 2002; Zhao and Chen, 2004].

Interaction systems often involve evolutionary dynamics. Several studies employed selection gradient to model evolving populations [Khibnik and Kondrashov, 1997; Mougi and Iwasa, 2010, 2011a,b], while others examined gene-for-gene coevolution [Sardanyes and Sole, 2006]. Khibnik and Kondrashov (1997) classified non-equilibrium (Red Queen) dynamics as evolution arising from fast ecological processes, slow genetic processes, or the combination of both (eco-evolutionary). The Red Queen hypothesis, which states that “species need to run or evolve in order to stay in the same place or to survive” [Van Valen, 1973; Rosenzweig et al., 1987], is often associated to host-parasite interaction. In this paper, we show illustrations presenting exceptional cases where evolutionary dynamics can explain the existence of oscillating population size.

We refer to the oscillating population sizes due to the structure of the constitutive interaction among species as *ecological oscillations*. While, we call the oscillatory behavior due to host-parasite coevolution as *evolutionary oscillations*.

## II. The Mathematical Model

We numerically investigate the behavior of host-parasite interaction system using an ODE model where there are m hosts and n parasites. This model does not only consider antagonistic interaction between hosts and parasites but also the inter-host and inter-parasite competition. We consider the following general (multispecies) model:

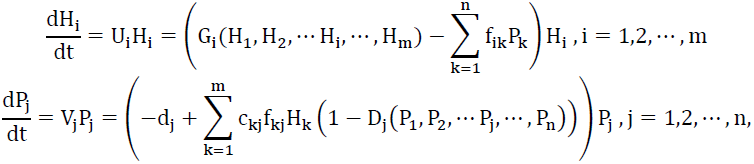

where H_i_ and P _j_ respectively denote the population of host i and parasite j. The parameters U_i_ and V_j_ are the corresponding fitness functions. Refer to Ref. Appendix I for the description of the variables and parameters.

The change in the size of host population is positively influenced by its growth rate G_i_(H_1_, H_2_,⋯ H_i_,⋯, H_m_), which is affected by the basal per capita birth rate and possibly by inter-host competition. However, it is negatively influenced by parasitism. In the absence of parasitism, host populations never reach extinction. One parasite can exploit a fraction of the host population by a factor dictated by the functional response. On the other hand, the parasite population is positively influenced by the reproduction rate of the parasite due to its utilization of hosts (numerical response), and negatively influenced by the death rate and competition among parasite species. In the absence of hosts, all parasite populations vanish.

There are two common mathematical representations of the host growth terms, namely, model with constant effective growth rate (exponential or the Malthusian), and model involving inter-host competition and carrying capacity (logistic). In the absence of parasitism, a host population with constant effective growth rate will exponentially propagate without bound. Whereas, a host population with growth rate that is affected by the carrying capacity and competition among hosts approaches equilibrium. Similarly, the parasite growth rate can involve inter-parasite competition. The term D_j_ refers to the situation where parasite populations can be satiated by other resources other than the hosts.

### Coevolutionary Dynamics

We consider cases where there are coevolving species due to competition. In this paper, evolution is represented by a system involving the concept of quantitative traits (genetic, phenotypic or behavioral traits) and selection gradient. Let U_i_ and V_i_ be functions of host and parasite populations as well as of the values of the quantitative traits (e.g., traits that are related to the parameters in the functional response). We define u_ij_ as the mean quantitative trait of the i-th host population specific for dealing with the j-th parasite population. We similarly define v_ij_ as the mean quantitative trait of the j-th parasite population specific for dealing with the i-th host population. We suppose slow constant genetical changes represented by small values of the speeds of evolutionary adaptation (i.e., ε_ij_ ≪ 1 and δ_ij_ ≪ 1 respectively for the host and parasite population). The representation of the coevolutionary dynamics is as follows:

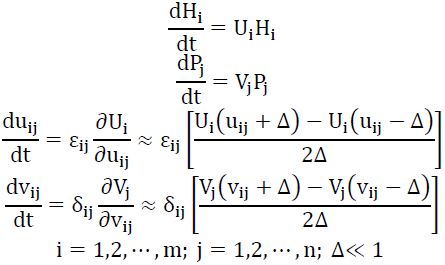

A positive selection gradient (value of the partial derivative) drives the population to climb a stronger trait value, and a negative gradient drives the population to have a lower trait value. For example, when U_i_ decreases due to the increase in u_ij_, then the value of u_ij_ should be reduced for the benefit of the host species. Alternatively, when U_i_ increases due to the increase in u_ij_, then the value of u_ij_ should be improved. This scenario shows that the host develops defense to counteract the parasite. Sexual reproduction and biological diversity often play big roles in this kind of evolutionary process. On the other hand, the parasite also evolves to increase or decrease the value of v_ij_ in response to the host’s evolution. Several studies have shown various empirical evidences of temporal coevolutionary dynamics [Decaestecker et al., 2013].

Progressive evolution has a trade-off since evolution entails costs and an indefinitely advancing trait is unlikely. In this paper, a climb from an inferior trait to a stronger trait results to a decline in the birth rate of the evolving population. For example, the host’s growth rate can be represented by a rational function 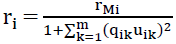 to characterize the evolutionary trade-off. The parameter r_Mi_ is the positive basal (maximal) birth rate and each q_ij_ denotes the degree of trade-off that affects the shape of the trade-off function curve. The rational trade-off functions assure that the value of r_i_ is always in the interval [0, r_Mi_] for any positive trait value.

Unlike most models where only one host and one parasite are involved, multispecies interaction are frequently asymmetric [Dawkins and Krebs, 1979; Lapchin and Guillemaud, 2005]. Parasites can select their host, while hosts do not choose their parasite. Our model can accommodate this situation by having asymmetric interaction parameter values. Evolving quantitative traits can also be asymmetric.

## III. Interaction Without Carrying Capacity and Without Coevolution

The exponential and logistic growth terms become approximately equivalent when the carrying capacity factor K for the host populations is very large 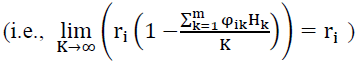. Correspondingly, when the carrying capacity 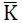 for the parasite populations is very large, the limiting term affecting the growth rate of the parasite becomes approximately zero (D_j_ ≈ 0). For these reasons, we give more emphasis on models with carrying capacities (both in host and parasite populations), since in nature, a population cannot grow unboundedly as t → ∞ even in the absence of competition. Nevertheless, as a take-off point, let us first discuss the Lotka-Volterra (LV) model without considering any carrying capacity (or with infinite resources).

The following Lotka-Volterra system with exponential growth (for host) and decay (for parasite) terms, and Holling type I functional response

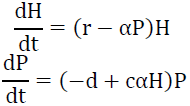

has two equilibrium points (namely, (0,0) and 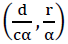), which are unstable. However, the solution (given any nonnegative initial condition and parameter values) is bounded and is actually always approaching a stable limit cycle. This model is structurally unstable [Murray, 2002] but it can be used as a groundwork for a more realistic representation.

In the general LV system 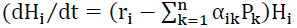, i = 1,2,…,m; 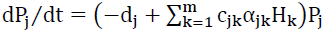, j = 1,2,…,n), the situation where all host and parasite populations are extinct is unstable. However, unlike single host-single parasite LV system where species always coexist, it is possible in the general system for some but not all host and parasite populations to vanish (i.e., populations may go extinct but at least one host and one parasite survive).

The surviving populations have higher fitness (e.g., high value of birth rate or low value of death rate) compared to the extinct species. Moreover, the amplitude of the solution corresponding to the surviving population can be greater than the other surviving species depending on the initial condition. The dynamics of this multispecies interaction implies that the superior populations are more progressive; and the weakest remains inferior or possibly, verge to vanish.

A small perturbation in the birth rate of a host species or in the death rate of a parasite causes bifurcation, specifically when all initial conditions and parameter values are equal. A small difference in the birth rates of hosts (or in the death rates of parasites) induce extinction of inferior species. It is possible that only two species will survive in the long run (i.e., the superior host with the highest birth rate and the superior parasite with the lowest death rate remain) which reduces the general LV model to a single host-single parasite system (see Figure 1).

**Figure 1.**
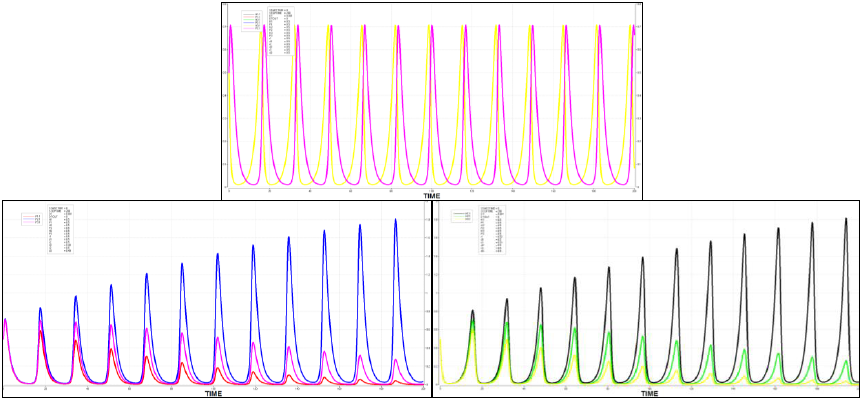
Influence of small change in birth rate of host and death rate of parasite may induce extinction in populations.

The solution to the general LV system always oscillates (ecological oscillations). Murray (2002) discussed insights regarding possible instability in a multispecies LV system and the danger of naively fitting models without biologically sound foundations to explain oscillating empirical data. A small change in the initial condition can produce large phase shifts in the solution after several time steps. At some periods, the population density of a surviving population declines too much near the edge of extinction because of very relatively large amplitude. These risky phenomena are common in oscillating systems.

Moreover, Figure 2 shows perturbation of a parameter value that results to a bifurcation from seemingly simple oscillatory behavior to a peculiar oscillation. In the classical LV model, chaos is not possible, but in a general model (with three or more species), chaos may arise. The occurrence of strange attractors prompts potential unpredictability and inadequacy of the LV model in dealing with biological data. When performing parameter estimation (fitting experimental data), it is advisable to examine if the solution to the fitted LV model is not sensitive to small perturbations, such as measurement errors.

**Figure 2.**
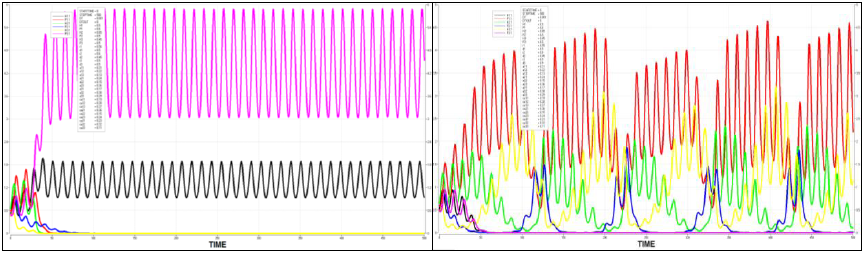
Influence of small change in parameter value (in this case, r_1_ = 0.56 to r_1_ = 0.55) results to a shift in behavior – from a seemingly simple oscillation to a peculiar one.

**Figure 3.**
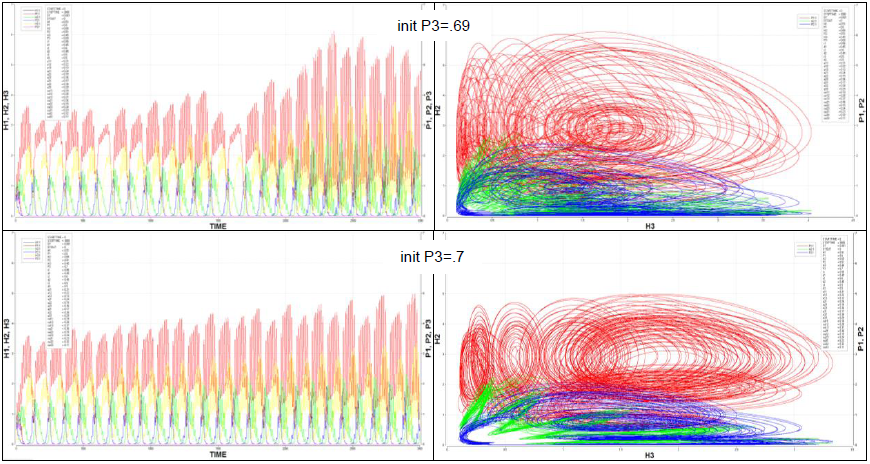
Multispecies Lotka-Volterra model appears to exhibit chaos given certain parameter values. Sensitivity to initial conditions can be observed (in this case, P_3_(0) = 0.69 to P_3_(0) = 0.7).

## IV. Interaction With Carrying Capacity but Without Coevolution

We investigate the solution to the following modified Rosenzweig-MacArthur (RM) ODE model:

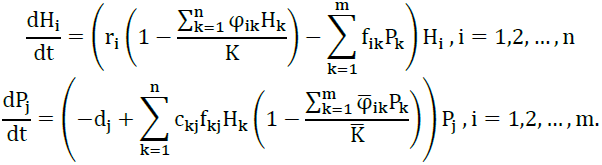

The host and parasite species with the greatest fitness respectively dominate the other host and parasite populations. Unlike LV model, populations in the RM model never propagate unboundedly and the size of any host (as well as any parasite) population never exceeds the carrying capacity. The solution to the RM model may approach an equilibrium point depending on the initial condition and parameter values.

The RM model has (0,0,⋯,0) as an equilibrium point, denoting a system where all species are extinct. However, this point is unstable. It follows that when there is a host population with nonzero initial condition then we expect that at least one host species survives. Although, if the death rates of all parasite populations are higher than the growth rates generated by utilizing host populations then all parasite population can vanish.

The equilibrium point (H*, P*) = (h, 0) such that h > 0 in single host-single parasite system is always stable from the positive value of P. For any given functional response, such equilibrium point always has h = K/φ. Convergence to this equilibrium point implies that the host survives while the parasite becomes extinct. This phenomenon is not surprising since when the parasite vanishes, the model reduces as a classical single species Verhulstian (logistic) model where host population converges to K/φ (where K is the host’s carrying capacity). Moreover, it is impossible to have an equilibrium point (H*, P*) = (0, p) such that p > 0. This portrays our assumption that the parasite population cannot survive without a host.

Many computational and mathematical studies have been done involving two (classical) as well as three (one host and two parasites, or two hosts and one parasite) species interaction, and some employed the classical model as part of a hybrid model. However, as more state variables (representing population of species) and parameters are added, the model becomes more challenging to analyze.

Furthermore, in a system with two or more host species (or with two or more parasite species), infinitely many equilibrium points may exist, specifically in a model with Holling type I where two species have the same parameter values. This implies that it is possible for two species with very similar characteristics to have different future population sizes when their initial populations are not the same.

In the RM model with Holling type I functional response (see Figure 4), larger carrying capacities both for host and parasite populations induce apparent ecological oscillation in population sizes. However, the oscillating solution does not necessarily converge to a limit cycle (it converges to a stable focus), unless, for instance, when carrying capacity is infinite. In addition, the host and parasite populations do not necessarily saturate the carrying capacity as opposed to the classical single species Verhulstian (logistic) model. In some situations, every population size for host or parasite species is less than the carrying capacity as a result of parasitism and interspecies competition.

Long-term apparent oscillations always occur when host and parasite interaction has appropriate parasitism functional response (e.g., Holling Type I) coupled with unbounded growth rates (immense carrying capacity). This phenomenon is related to the Paradox of Enrichment. Boundless growth potential and parasitism efficiency enable the populations of the interacting species to reach extreme states.

Ecological oscillations associated with a large carrying capacity are commonly fostered by certain functional responses, such as Holling type I, where there is unrestrained (no satiation) parasite utilization efficiency. If other functional response is used, it is possible that a large carrying capacity does not necessarily result to long-term apparent oscillation. Functional response plays a big role in producing oscillatory behavior. Note: oscillations can also arise given Holling type II and III for some parameter values. In this paper, we focus on Holling type I.

**Figure 4.**
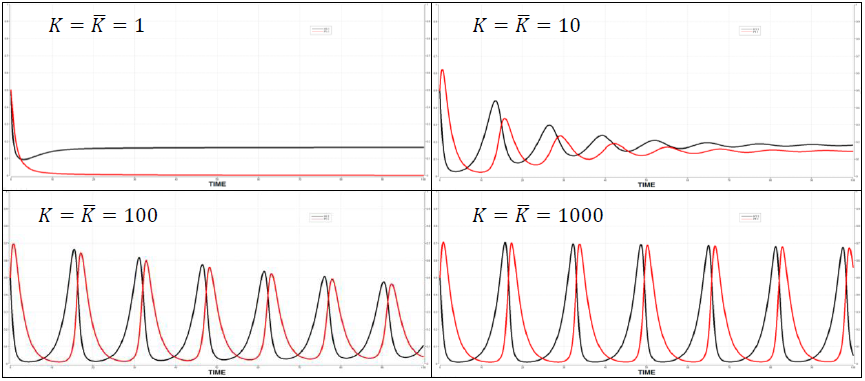
A larger carrying capacity induces oscillation.

Furthermore, similar to LV models, RM models with Holling type I functional response may result to peculiar ecological oscillations, especially when carrying capacities are large. Some of these peculiar fluctuations in the trajectory of the ODE show sign of chaos-like behavior.

Long-term apparent oscillations are not only possible because of large carrying capacity. There are cases that changing the birth and death rates as well as initial conditions of the host and parasite species result to simple or peculiar ecological oscillations. There are also cases where ecological oscillations occur depending on the structure of parasitism efficiency matrix. A right combination of parameter values can generate peculiar oscillatory solution (see Figure 5).

**Figure 5.**
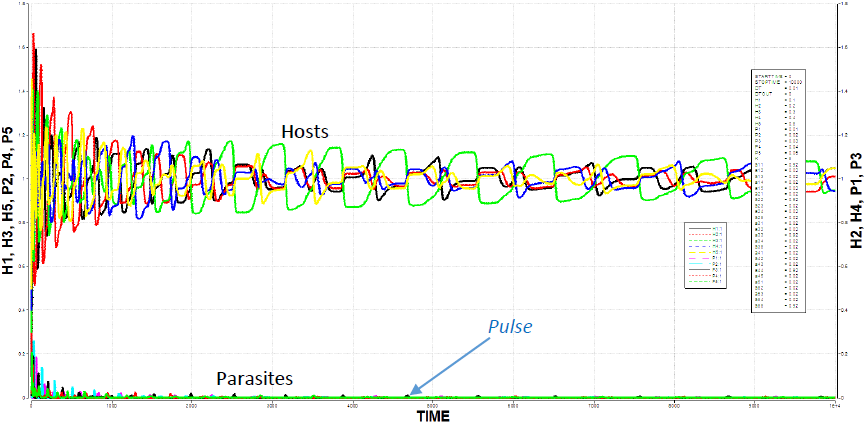
Certain parameter values can generate long-term apparent oscillatory behavior.

Figure 5 presents a case where parasitism efficiency matrix is of the form

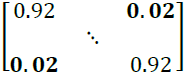

This matrix shows that *H*_*i*_ is the main host of parasite *i*. While all *H*_*k≠i*_ are alternative hosts. In this example, parasite populations approach the edge of extinction but they create pulses that induce oscillations in the population of the hosts.

## V. Interaction With Carrying Capacity and With Coevolution

Quantitative traits and population densities of the interacting species are affected by evolution. For example, suppose there are 2 identical host populations (H_1_ and H_2_) and 2 identical parasite populations (P_1_ and P_2_). If H_1_ evolves against P_1_, it is possible that H_1_ will dominate H_2_, and P_2_ will dominate P_1_. According to the Red Queen Hypothesis, the parasites and hosts must coevolve to counteract their evolutionary disadvantage. However, note that evolution does not necessarily result to survival.

Oscillating population sizes are possibly due to the constitutive interaction between species and not because of coevolution. The oscillatory behavior of interacting host and parasite species, without looking at the evolving quantitative traits, are not enough evidence of the Red Queen. We need to look at evolutionary oscillations, which are beyond the ecological oscillations. To establish this claim, we present illustrations of the interaction among the host and parasites, with and without coevolutionary dynamics. Figure 6 shows that oscillations are not easily generated by the Red Queen.

**Figure 6.**
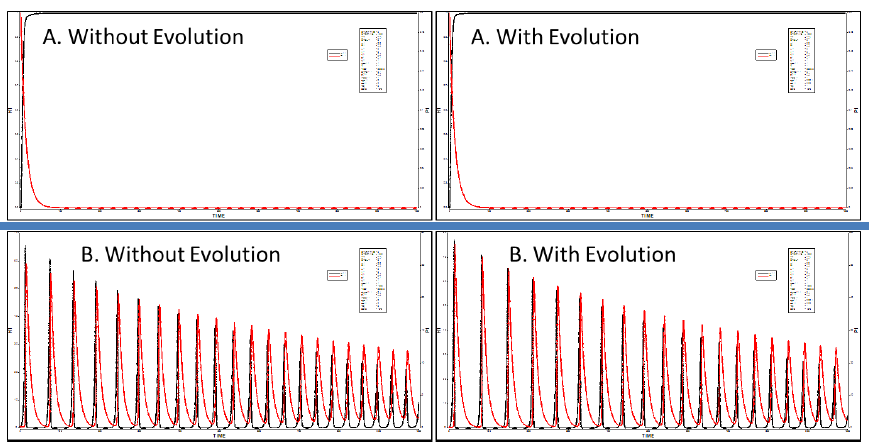
Oscillations are not necessarily evolutionary. Oscillations in the system with coevolutionary dynamics are inherited from the constitutive interaction among species.

Coevolution does not always result to long-term apparent oscillating population size (beyond the oscillations caused by constitutive interaction). Figure 7 presents examples of the exceptional cases where the Red Queen generates oscillation. We can observe in Figure 7 the difference between evolutionary oscillation and ecological oscillations. Evolutionary oscillation is seen by looking at evolutionary time scale. Although, a non-evolving species (e.g., parasite) may seemingly exhibit evolutionary oscillations because other competing species (e.g., other parasites) are evolving. In this case, evolutionary oscillation turns to be ecological, too. In addition, the speed of evolutionary adaptation can dictate the existence and the period of the apparent oscillations.

**Figure 7.**
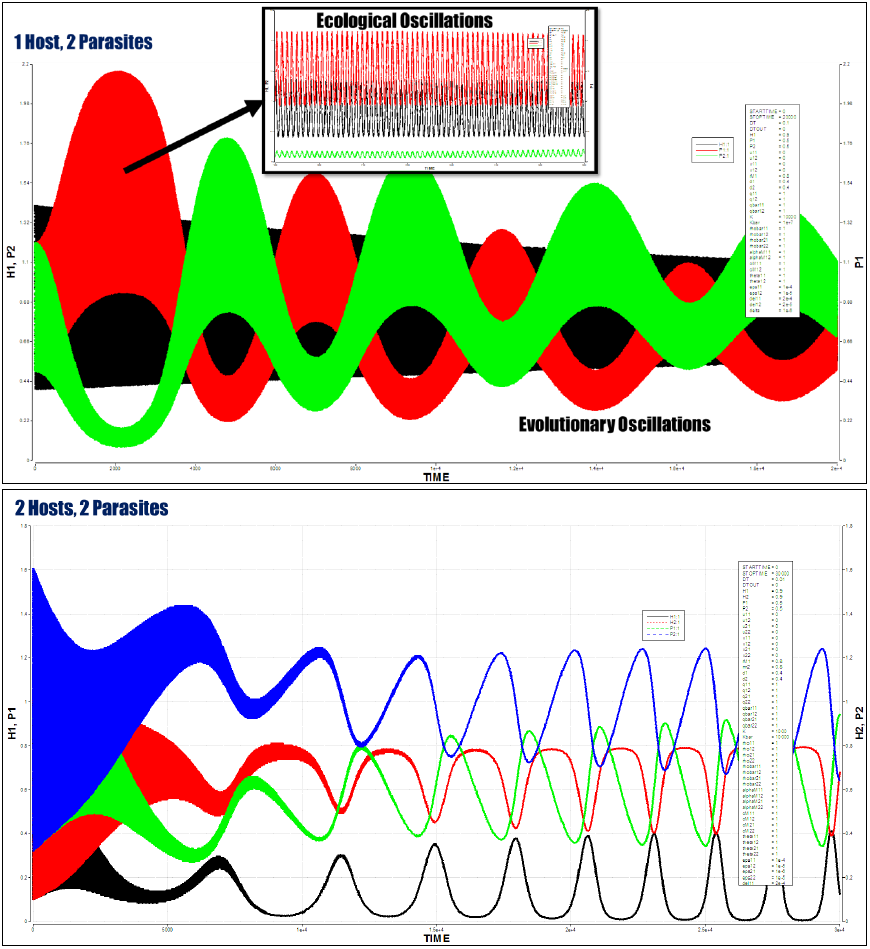
Evolutionary oscillations observed at evolutionary time scale beyond the ecological oscillations.

The illustrations shown in Figure 7 are oscillations that became possible with the aid of inter-parasite competition. We hypothesize that in our ODE system, long-term apparent evolutionary oscillations are not possible in single host, single parasite interaction. The Red Queen reveal herself through oscillating population sizes when there are three or more interacting species (inter-parasite as well as inter-host). However, we do not dismiss the case where evolution affects ecological oscillations. Sometimes ecological and evolutionary oscillations are mixed and indistinguishable from each other, especially when the evolving traits push the system to have a set of parameter values that results to ecological oscillations or when there are already existing peculiar ecological oscillations (see Figure 8).

**Figure 8.**
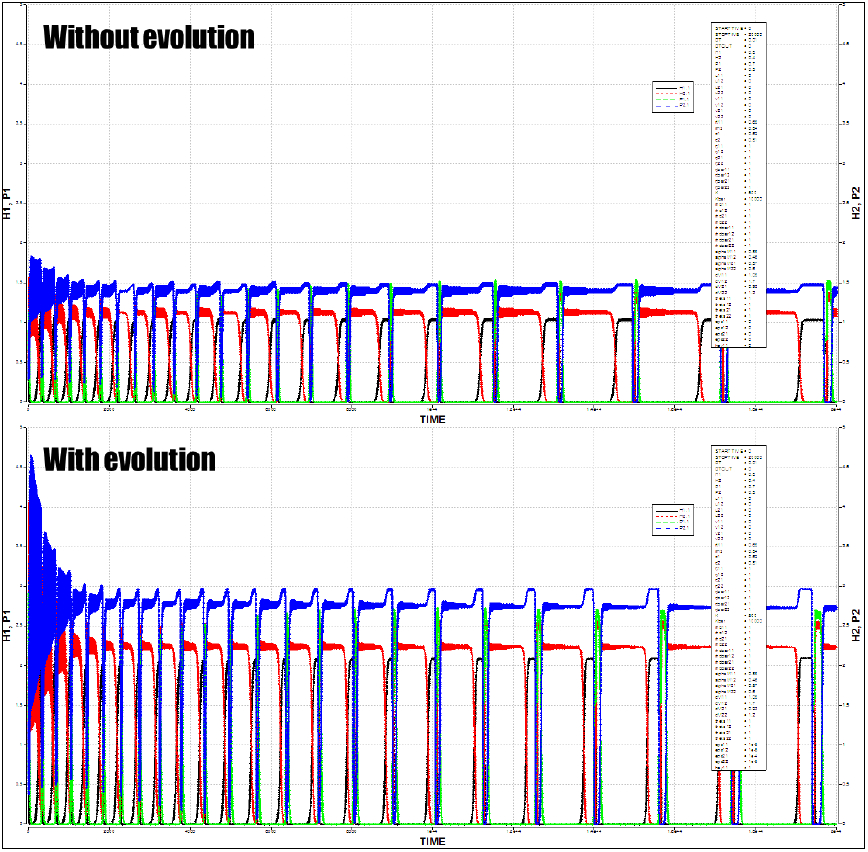
An example where there are already peculiar ecological oscillations, and evolution affects the behavior of the ecological oscillations (e.g., change in amplitude).

A right combination of parameter values can generate oscillations. Speed of evolutionary adaptation and degree of trade-off play significant roles in generating evolutionary oscillation in population size. Low and high values of trade-off parameters do not provide good conditions for oscillatory behavior. In fact, a high value of trade-off parameter may constrain the evolution of quantitative traits. However, introducing some medium-level degree of trade-off during co-evolution results to oscillations (see Figure 9 for an example).

**Figure 9.**
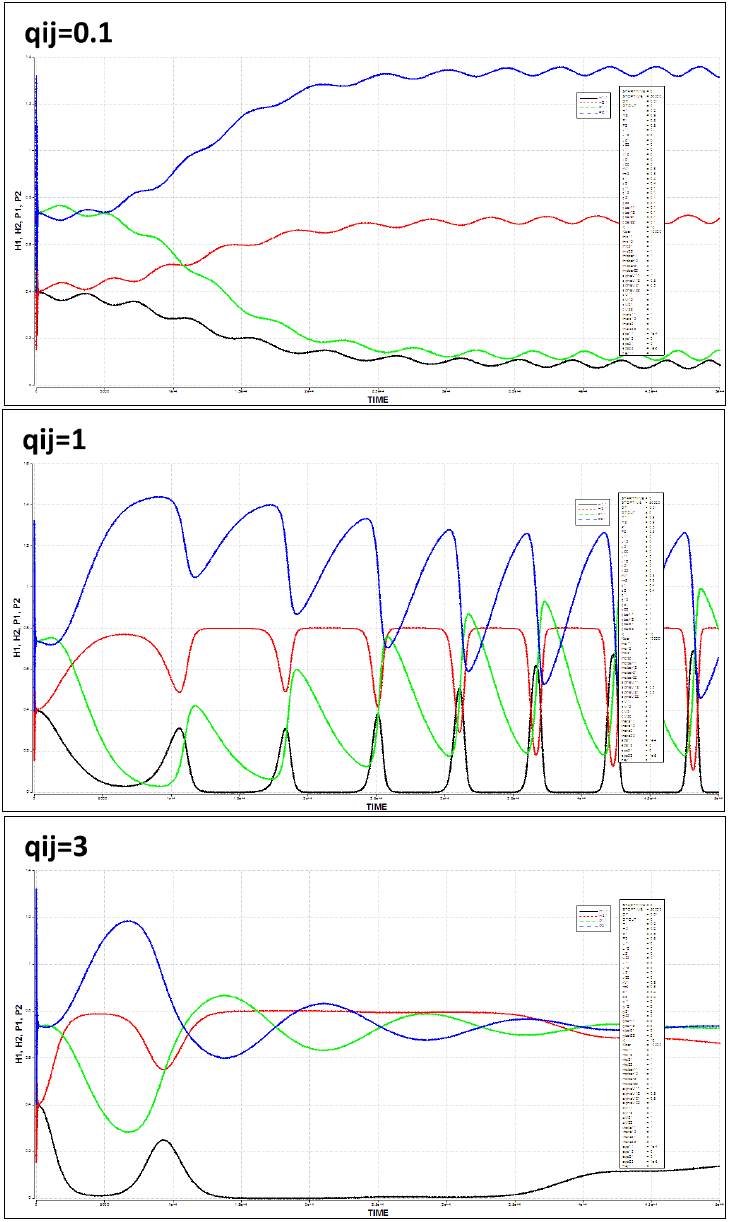
Evolutionary trade-off does not always result to oscillatory behavior. There is a range of parameter values that generates evolutionary oscillations.

In a system without coevolution, the equilibrium values depend on the parameters. Thus, if parameters become dynamic because of evolution then it is possible that the trajectory of the ODE solution shifts equilibrium values, which results to evolutionary oscillations. However, there are some cases when coevolution shutdowns ecological oscillations.

In our model of coevolutionary dynamics, the fitness functions (as well as the population densities) and the quantitative traits operate in a feedback loop. Evolution of the quantitative traits affects the population densities through the fitness functions; and the fitness functions dictate the evolution of the traits. The Red Queen hypothesis indicates that coevolving species undergo endless arms race competition (e.g., unbounded set of feasible phenotypes) [Rosenzweig et al. 1987; Dieckmann et al. 1995], while balancing the benefit and cost of evolution. There are cases when coevolution exhibits unbounded non-equilibrium dynamics depicted in the quantitative traits. Some argue that this unbounded dynamics is rare or possibly unrealistic [Dieckmann et al. 1995]. However, there are cases when evolving quantitative traits tend to an attractor (equilibrium point, limit cycle or strange attractor). Arms race competition can end [Dawkins and Krebs, 1979].

Figure 10 presents an example of evolving quantitative traits of hosts and parasites. The quantitative traits of the hosts approach an attractor but the quantitative traits of the parasites continuously increase. Pressure is given to the parasite species since parasite species can vanish if it cannot run after the evolving host, yet it cannot kill all the hosts to avoid extinction (since parasites cannot live without a host). This example is one of the numerous scenarios that arise from coevolutionary dynamics. There are cases when coevolution results to extinction of one of the evolving host species. There are also cases when cryptic dynamics occur.

**Figure 10.**
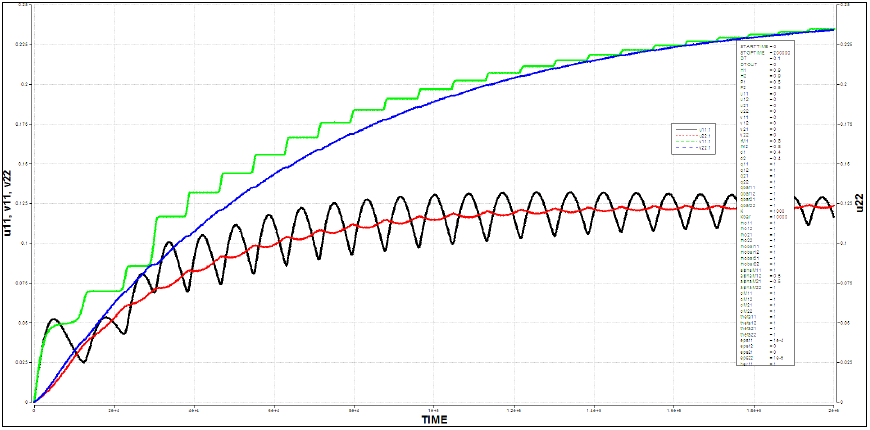
An example of evolving quantitative traits during coevolution.

## VI. Concluding Remarks

We numerically investigated several cases yielding to oscillating host-parasite populations, and we found that the Red Queen hypothesis can explain some of the exceptional cases (evolutionary oscillation). Arms race competition can be observed by tracking the evolution of quantitative traits, and it is possible that evolutionary oscillations can be recognized beyond ecological oscillations. One limitation of our model is that it cannot track polymorphism and the diversity of the changing phenotypes over the fitness landscape.

There are many possible model representations for the host-parasite or prey-predator system, justified using either mechanistic or empirical explanations. To have biological relevance, we should always ensure that for any finite time, a unique solution to the model exists and that state variables should always be non-negative. Parameter values can be estimated using several techniques, such as by curve fitting or by machine learning. Keen investigation is important in determining the robustness of the model and if empirical observations match the behavior of the theoretical model to be used in representing biological phenomena.

Ecological and evolutionary oscillations result from a right combination of parameter values. This study can be extended by considering other functional response curves, non-monotonic evolution function and spatial distribution of species. Demographic and environmental randomness can also be incorporated to determine if the oscillations are robust against stochastic noise.

## APPENDIX I Definition of Variables and Parameters

APPENDIX I: Definition of Variables and Parameters
| State Variables and Parameters | Definition |
| --- | --- |
| $m$ | number of different host species |
| $n$ | number of different parasite species |
| $H_i$ | population of host $i$ |
| $P_j$ | population of parasite $j$ |
| $U_i$ | host fitness function |
| $V_j$ | parasite fitness function |
| $G_i(H_1, H, \dots H_i, \dots, H_m)$ | effective growth rate of the population of host $i$; Exponential:<br>$G_i(H_1, H, \dots H_i, \dots, H_m) = r_i$ or Logistic:<br>$G_i(H_1, H, \dots H_i, \dots, H_m) = r_i \left(1 - \frac{\sum_{k=1}^m \varphi_{ik} H_k}{K}\right)$ |
| $r_i$ | basal growth rate of the population of host $i$ |
| $\varphi_{ik}$ | nonnegative relative strength of host $i$ over host $k$ |
| $K$ | size of the carrying capacity of the environment of host populations during inter-host competition |
| $D_j(P_1, P_2, \dots P_j, \dots, P_n)$ | limiting term affecting the growth rate of parasite $j$ (e.g., due to inter-parasite competition for resources other than the host); suppose $D_j(P_1, P_2, \dots P_j, \dots, P_n) = \frac{\sum_{k=1}^m \bar{\varphi}_{jk} P_k}{\bar{K}}$ |
| $\bar{\varphi}_{ik}$ | nonnegative relative strength of parasite $j$ over parasite $k$ |
| $\bar{K}$ | size of the carrying capacity of the environment of the parasite populations during inter-parasite competition. This carrying capacity includes resources excluding host utilization. |
| $d_j$ | constant death rate of the population of parasite $j$ |
| $f_{ij}$ | the parasitic utilization efficiency (part of the functional response) |
| $c_{ij}$ | conversion rate or birth rate of parasite due to host utilization |
| $u_{ij}$ | mean quantitative trait of the $i$-th host population specific for dealing with the $j$-th parasite population |
| $v_{ij}$ | mean quantitative trait of the $j$-th parasite population specific for dealing with the $i$-th host population |
| $\varepsilon_{ij}$ and $\delta_{ij}$ | speeds of evolutionary adaptation of the host and parasite populations, which are functions of the additive genetic variances |

## APPENDIX II Functional Response Curve

Let *F*_*ij*_ = *f*_*ij*_*H*_*i*_. We consider the following monotonic one-variable functional response curve:

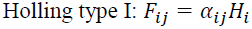

where the nonnegative constant parameter *α*_*ij*_ is defined as the efficiency of the parasite in utilizing hosts (i.e., one parasite can parasitize *α*_*ij*_*H*_*i*_ number of hosts). The matrix containing the *α*_*ij*_ ’s is called the parasitism efficiency matrix [*α*_*jy*_].

The Holling type I functional response represents linear curve, and only depends on the size of host population [Piana et al. 2006; Kratina et al. 2009]. Holling type 1 indicates that the rate of utilization of one parasite is directly proportional to the size of host population, where the ratio between the number of parasitized hosts and the parasite population is a linear function 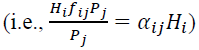.

## APPENDIX III The Selection Gradient

We also consider cases where there are co-evolving species due to competition. In this paper, evolution is represented by a system involving the concept of quantitative traits (genetic, phenotypic or behavioral traits) and selection gradient [Khibnik and Kondrashov, 1997; Mougi and Iwasa, 2011a,b]. Let *U*_*i*_ and *V*_*i*_ be functions of host and parasite populations as well as of the values of the quantitative traits (e.g., traits that are related to the parameter *α*_*ij*_). In this paper, We assume that the evolving quantitative traits are only those related to *α*_*ij*_. We define *u*_*ij*_ as the mean quantitative trait of the *i*-th host population specific for dealing with the *j*-th parasite population. We similarly define *v*_*ij*_ as the mean quantitative trait of the *j*-th parasite population specific for dealing with the *i*-th host population. We suppose slow constant genetical changes represented by small values of the speeds of evolutionary adaptation (which are actually functions of the additive genetic variances), that is, *ε_ij_* ≪ 1 and *δ_ij_* ≪ 1 respectively for the host and parasite population. A positive selection gradient (value of the partial derivative) drives the population to climb a stronger trait value, and a negative gradient drives the population to have a lower trait value. For example, when *u*_*i*_ decreases due to the increase in *u*_*ij*_, then the value of *U*_*ij*_ should be reduced for the benefit of the species. On the other hand, when *U*_*i*_ increases due to the increase in *u*_*ij*_, then the value of *u*_*ij*_ should be improved for the benefit of the species. The representation is as follows:

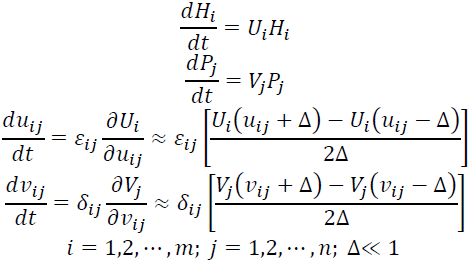

The evolving parameters (such as *α*_*ij*_) due to the evolving quantitative traits can be modeled using a monotonic curve (e.g., 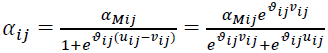 which is sigmoidal) [Mougi and Iwasa, 2011a,b]. The value of *ϑ*_*ij*_ ≥ 0 defines the steepness of the curve, and the parameter *α*_*Mij*_ is the basal and possible maximal value of *α*_*ij*_. Monotonic curves are usually used to represent co-evolution due to arms-race competition.

Progressive evolution has a trade-off since evolution entails costs and an indefinitely advancing trait is unlikely [Khibnik and Kondrashov, 1997]. In this paper, a climb from an inferior trait to a stronger trait results to a decline in the birth rate (e.g., *r*_*i*_ and *c*_*ij*_) of the evolving population. Let 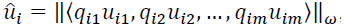, and for simplicity, we assume ω = 2 (2-norm), that is, 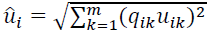. The parameter *û*_*i*_ represents the collective trait of the host population, where the *q*_*ij*_’s are parameters that affect the shape of the trade-off function curve. There are various representations of the trade-off function, such as the standard polynomial function 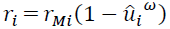 [Mougi and Iwasa, 2011a,b]. However, we suppose the effect of evolution to the birth rates are represented as rational functions instead of polynomials such as 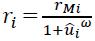 and 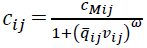 where *r*_*Mi*_ and *c*_*Mij*_ are positive basal (maximal) birth rates, and the exponent (ω ≥ 2) as well as 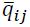 are parameters that define the shape of the curves. Similar to the usual polynomial functions, if the value of the exponent ω = 1, then the system may have a negative trait value which is unrealistic [Mougi and Iwasa, 2011b]. In contrast to the usual polynomial functions, the rational trade-off functions assure that the values of *r*_*i*_ and *c*_*ij*_ are always in the interval [0, *r*_*Mi*_] and [0, *c*_*Mij*_], respectively, for any positive trait value.

